# ABCC10 roles in plant development and the transport of indole-3-butyric acid

**DOI:** 10.1101/2024.02.07.579300

**Authors:** Arielle L. Homayouni, Suresh Damodaran, Katherine Schreiber, Marta Michniewicz, Lauren K. Gunther, Lucia C. Strader

## Abstract

Proper spatiotemporal distribution of the phytohormone auxin throughout plant tissues mediates a variety of developmental processes. Auxin levels are tightly regulated via *de novo* synthesis, transport, and conversion from its conjugated forms and precursors. These levels can be regulated through conversion of the auxin precursor, indole 3-butyric acid (IBA), into the active auxin, indole-3-acetic acid (IAA), in a peroxisomal β-oxidation process. Defects in IBA-to-IAA conversion cause multiple developmental defects in Arabidopsis, demonstrating IBA-derived IAA is physiologically important to the active auxin pool. Similar to IAA, transport of IBA modulates development. However, the mechanisms governing transport of this molecule remain largely unknown. Here, we identify a mutation in the *ABCC10* gene of Arabidopsis that suppresses the *abcg36* hypersensitivity to IBA and its synthetic analog, 2,4-dichlorophenoxy butyric acid (2,4-DB) and the *abcg36* hyperaccumulation of [^3^H]-IBA. We found that ABCC10 acts as a direct vacuolar transporter of IBA. Further, *ABCC10* is necessary for proper development of the root apical meristem and leaf tissue. Our findings uncover a previously uncharacterized method of IBA transport that regulates aspects of plant development.

## INTRODUCTION

Plants modulate their development and metabolism in response to environmental stimuli. Their ability to alter the availability and distribution of phytohormones, such as auxin, is essential to these responses (reviewed in Santner and Estelle, 2009). Auxin regulates plant growth and development through its orchestration of cell division, elongation, and differentiation from embryogenesis through maturity (Perrot-Rechenmann, 2010; reviewed in Ljung, 2013). Proper spatiotemporal distribution of auxin within a plant is essential for many adaptive responses (reviewed in Vanneste and Friml, 2009) and is regulated on a cellular level to ensure precise organ morphogenesis and physiology. The appropriate level and distribution of active auxin (indole-3-acetic acid, IAA) is tightly regulated via *de novo* synthesis, transport, and conversion from its conjugated forms and precursors (reviewed in Korasick et al., 2013).

Transport and conversion of the auxin precursor indole-3-butyric acid (IBA) contributes to auxin homeostasis and plant developmental events (reviewed in Damodaran and Strader, 2019). IBA and IAA are virtually identical compounds, except IBA carries a four-carbon side chain whereas IAA carries a two-carbon side chain (Fig. 1A). IBA is converted to IAA through a multistep process that occurs in the peroxisome in a process similar to fatty acid β-oxidation (Zolman et al., 2000; Strader et al., 2010). Genetic evidence has revealed that IBA-to-IAA conversion contributes to the auxin pool to regulate multiple developmental programs (Frick and Strader, 2018). Blocking IBA contributions to the auxin pool cannot be compensated by other IAA input pathways (Strader and Bartel, 2011), suggesting the spatiotemporal distribution of IBA-derived IAA leads to physiologically important changes in the auxin homeostasis.

**Figure 1.**
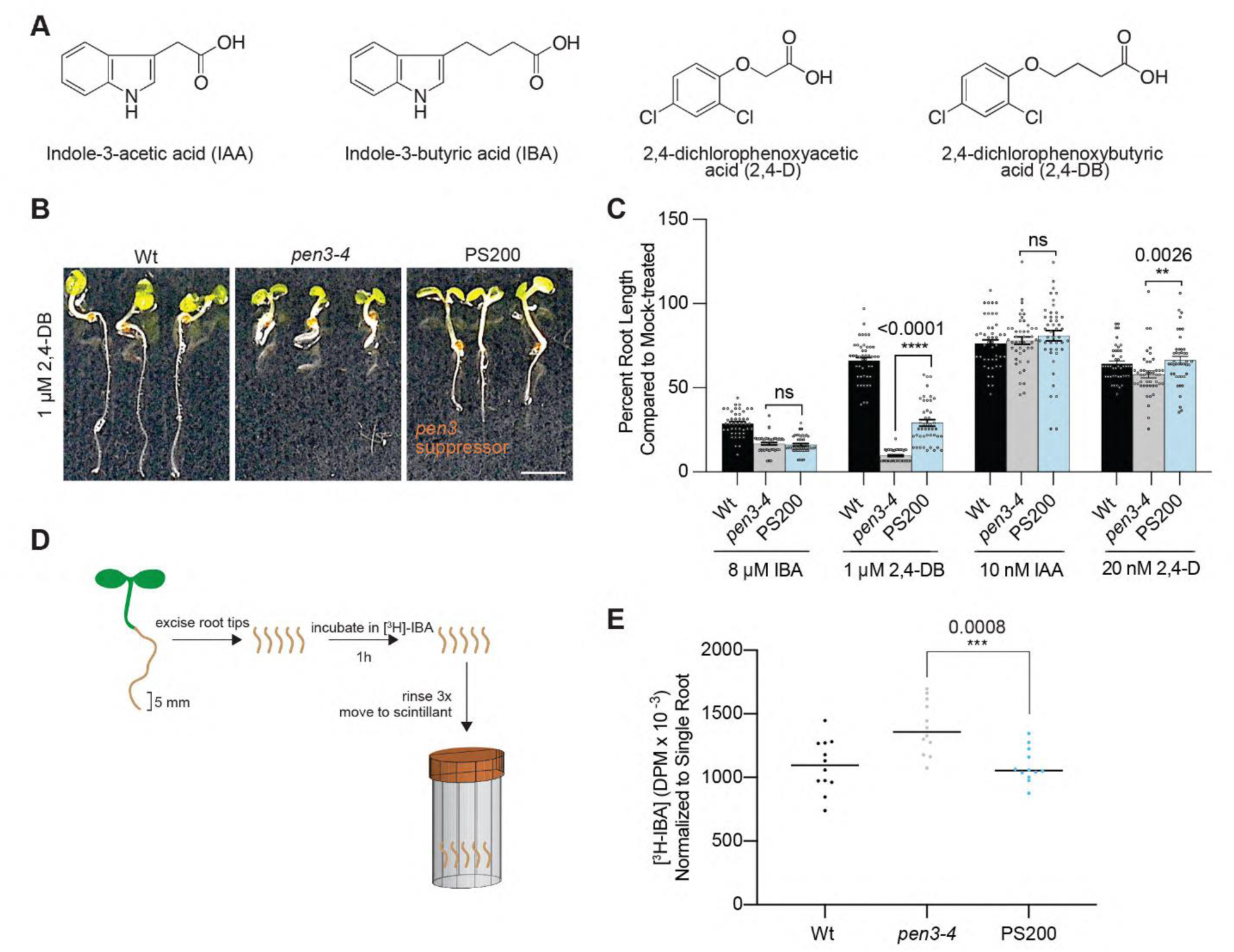
PS200 was isolated as a suppressor of the *abcg36* hypersensitivity to 2,4-DB. **A)** Chemical structure of indole-3-acetic acid (IAA) and indole-3-butyric acid (IBA), and their synthetic analogs 2,4-dichlorophenoxy butyric acid (2,4-DB) and 2,4-dichlorophenoxyacetic acid (2,4-D). **B)** Photograph of representative 7-day-old Wt (Col-0), *pen3-4*, and PS200 (*pen3-4 abcc10-*1) seedlings grown in the presence of 1 μM 2,4-DB. Scale bar, 5 mm. **C)** Mean normalized primary root lengths (+SEM; n ≥ 45) of 7-day-old Wt (Col-0), *pen3-4*, and PS200 (*pen3-4 abcc10-*1) seedlings grown in the presence of the indicated auxins and auxin precursors. F-values obtained from two-way ANOVA showed variation, so multiple comparisons using Dunnett’s test was perform with adjusted p-values indicated (∗p ≤ 0.05; ns, no significant difference observed). **D)** Schematic of root tip auxin transport assays in Arabidopsis. 5 mm seedling root tips are excised and incubated for 1 h in uptake buffer containing 25 nM [^3^H]-IBA, rinsed three times with uptake buffer, and then removed and analyzed by scintillation counting. **E)** Root tips of Wt (Col-0), *pen3-4*, and PS200 (*pen3-4 abcc10-*1) seedlings were used in auxin transport assays. Mean DPM normalized to root number, with 12 replicates of five root tips of each genotype, are shown. F-values obtained from one-way ANOVA showed variation, so multiple comparisons using Dunnett’s test was perform with adjusted p-values indicated (∗p ≤ 0.05).

Similar to IAA, IBA and IBA conjugates are transported in plants, yet many IBA carriers remain to be identified. Despite their near identical chemical structures, plants use distinct carriers to move IBA and IAA (reviewed in Damodaran and Strader, 2019). Several members of the Pleiotropic Drug Resistance Protein (PDR/ABCG) subfamily of ATP-binding cassette (ABC) transporters have been implicated in IBA efflux from root cells (Strader et al., 2008; Strader and Bartel, 2009; Růžička et al., 2010; Aryal et al., 2019). Specifically, both *ABCG36/PDR8/PEN3* and *ABCG37/PDR9/PIS1* are polarly localized to the outer face of root epidermal and lateral root cap cells, where they act to efflux IBA into the rhizosphere (Strader and Bartel, 2009; Růžička et al., 2010). Mutants defective in either *abcg36* or *abcg37* hyperaccumulate [^3^H]-IBA, leading to a hypersensitivity to both IBA and its synthetic analog, 2,4-dichlorophenoxy butyric acid (2,4-DB, Fig. 1A), but not IAA (Strader et al., 2008; Strader and Bartel, 2009; Růžička et al., 2010).

We have isolated and characterized a mutant that suppresses the hyperaccumulation of IBA in the *pen3-4* allele of *ABCG36*. Here, we report that loss-of-function mutations in *ATP-BINDING CASSETTE C10/MULTIDRUG RESISTANCE-ASSOCIATED PROTEIN 14* (*ABCC10/MRP14*) suppress both the *pen3-4* hypersensitivity to 2,4-DB and the *pen3-4* hyperaccumulation of IBA, phenotypes that are suppressed by mutants unable to uptake and/or transport IBA. We demonstrate that *abcc10* mutants in a wild-type (Wt) *ABCG36* background are mildly resistant to IBA and 2,4-DB and display developmental defects in root apical meristems and leaf expansion. Our characterization provides insight into transport of the auxin precursor IBA and is consistent with the possibility that *ABCC10* is an IBA transporter that acts to modulate IBA contributions to the active auxin pool to promote proper maintenance of apical root meristem length and leaf size.

## RESULTS

### ABCC10 is required for full IBA response in roots

Peroxisomal IBA-to-IAA conversion contributes to the active auxin pool to drive multiple developmental events (reviewed in Damodaran and Strader, 2019). Despite the significance of IBA contributions to auxin-regulated developmental processes, few mechanisms to regulate IBA inputs to the auxin pool have been identified (Michniewicz et al., 2019). Here, we describe a forward genetics approach used to identify factors that may act to modulate IBA levels.

Mutants defective in *ABCG36/PDR8/PEN3* display hypersensitivity to the auxin precursor IBA, but not to the active auxin IAA (Strader and Bartel, 2009), likely due to a defect in efflux of IBA from root cells (reviewed in Michniewicz et al., 2014). In addition, *abcg36* mutants hyperaccumulate [^3^H]-IBA in simplified auxin transport assays (Strader and Bartel, 2009), consistent with a defect in efflux of this molecule. To identify additional IBA transporters, we performed a suppressor screen of the *pen3-4* allele of *ABCG36* (Michniewicz et al., 2019). Isolate PS200 was identified from this screen because it displayed partial suppression of the *pen3-4* hypersensitivity to the inhibitory effects of 2,4-DB on root elongation (Fig. 1B and 1C). Isolate PS200 also displayed suppression of *pen3-4* hyperaccumulation of [^3^H]-IBA (Fig. 1D and 1E).

Using whole genome sequencing of bulk segregants (Thole and Strader, 2015), we identified six homozygous EMS-related mutations in PS200. One of these was a missense mutation in *At3g59140*, encoding a member of the MULTIDRUG RESISTANCE-ASSOCIATED PROTEIN/ATP-BINDING CASSETTE C (MRP/ABCC) subfamily of ABC transporters, MRP14/ABCC10 (Fig. 2A), hereafter called ABCC10, resulting in a Asp237-to-Asn substitution (Fig. 2A and 2B) in a conserved residue (Fig. 2C). ABC proteins use the energy of ATP binding and hydrolysis to translocate substrates across cellular membranes. There are 129 ABC proteins in Arabidopsis, of which several ABCBs have been implicated in auxin efflux (Sánchez-Fernández et al., 2001; reviewed in Borghi et al., 2019). ABCC proteins are full-length ABC transporters, many of which have been implicated in vacuolar transport of structurally diverse substrates in plants (Klein et al., 2006; Verrier et al., 2008). Arabidopsis has 14 functional ABCCs, each of which is conserved throughout the plant kingdom (Fig. 2D). Some AtABCC proteins, including AtABCC5, have been implicated in vacuolar sequestration of IAA or IAA-conjugates (Gaedeke et al., 2001). Thus, the molecular identity of *ABCC10* made it a compelling candidate for investigation as a putative IBA transporter; we named this missense allele *abcc10-1*.

**Figure 2.**
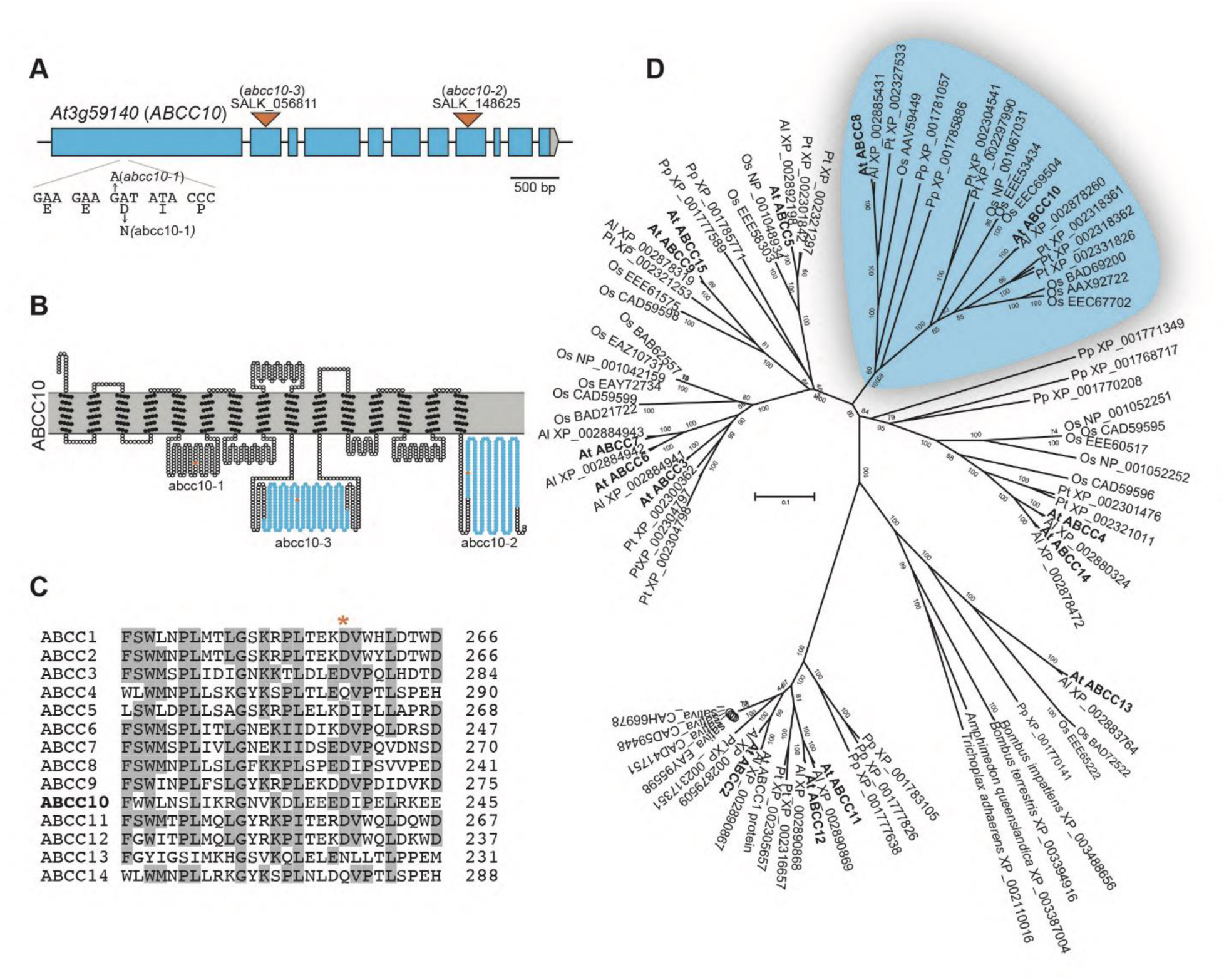
ABCC10 identification. **A)** *ABCC10/At3g59140* gene schematic depicting the EMS-consistent point mutation identified in PS200 (called *abcc10-1*) and two T-DNA insertion alleles (*abcc10-2* and *abcc10-3*). **B)** A schematic diagram of the predicted ABCC10 topology based on outputs of ARAMEMNON (http://aramemnon.uni-koeln.de; Schwacke et al., 2003) and TOPO2 (http://www.sacs.ucsf.edu/TOPO2/). Filled circles represent predicted transmembrane residues whereas open circles represent non-transmembrane residues. *abcc10-1* carries a Asp237-to-Asn substitution in a non-transmembrane domain, indicated by a filled red circle. The insertional T-DNA lines *abcc10-2* and *abcc10-3*, indicated by filled orange triangles, disrupt either of the predicted nucleotide binding domains, indicated by filled blue circles. **C)** The *abcc10-1* mutation disrupts a conserved acidic residue. The alignment shows this region in ABCC10 (At3g59140) and other members of the ABCC family in *Arabidopsis thaliana*. Because *AtABCC15* is a pseudogene (Kolukisaoglu et al., 2002), it was excluded from this alignment. **D)** Phylogenetic tree of ABCC protein sequences throughout the plant kingdom. The unrooted phylogram was generated using PAUP 4.05b (Wilgenbusch and Swofford, 2003) by performing the bootstrap method with 500 replicates. Bootstrap values are shown at the nodes.

We crossed our PS200 isolate (*abcg36-1 abcc10-1*) to wild type and obtained the *abcc10-1* mutant removed from the *abcg36-1* background. We also obtained two insertional alleles defective in *ABCC10*, which we named *abcc10-2* (SALK_148625) and *abcc10-3* (SALK_056811; Fig. 2A). Similar to *abcc10-1*, *abcc10-2* displayed mild resistance to IBA and 2,4-DB and wild-type sensitivity to the active auxins IAA and 2,4-dichlorophenoxyacetic acid (2,4-D; Fig. 3A and 3B), consistent with the possibility that the defect in *ABCC10* causes *pen3-4* suppression in isolate PS200.

**Figure 3.**
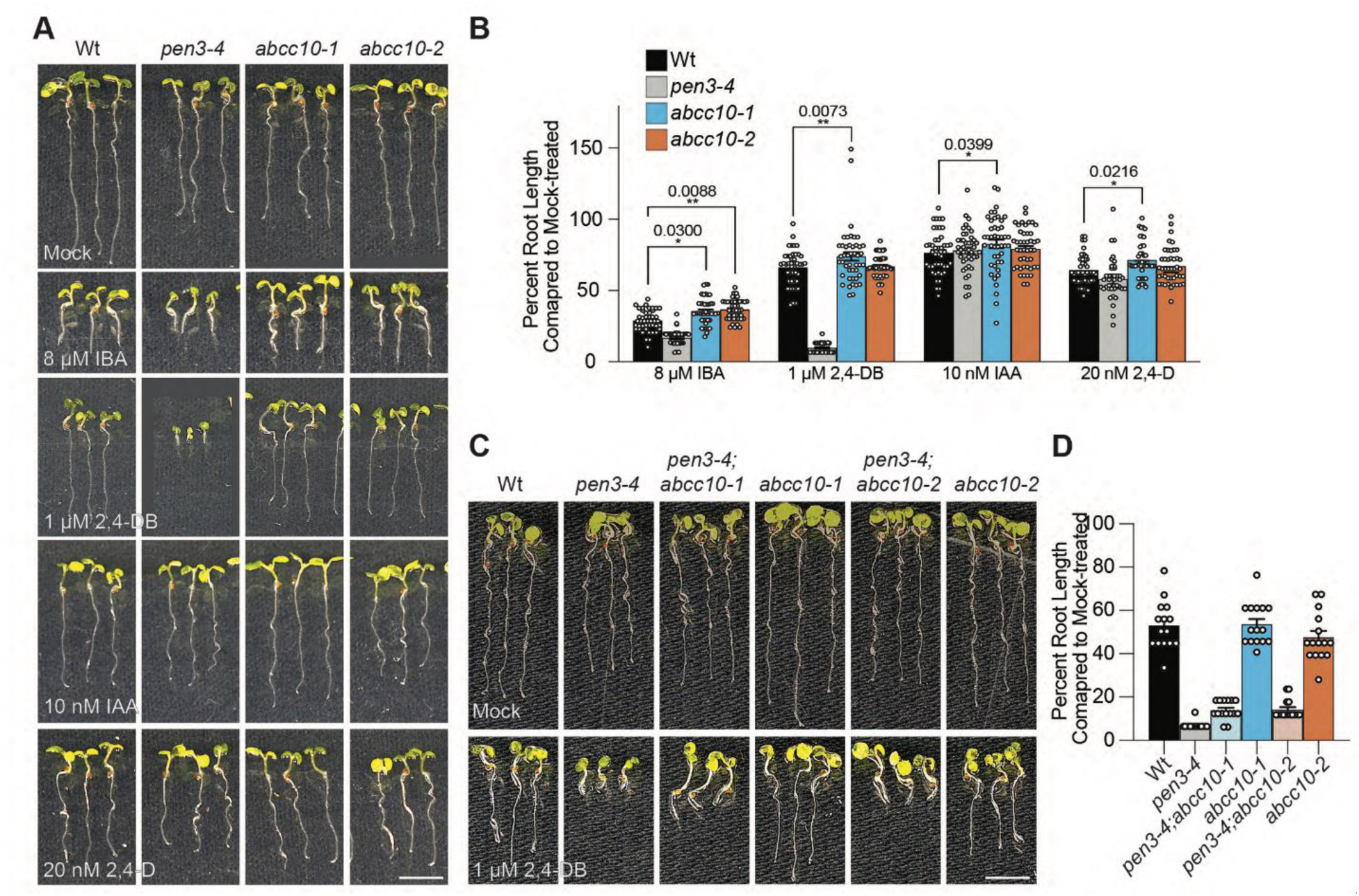
ABCC10 is required for full IBA response in roots. **A)** Photograph of representative 7-day-old Wt (Col-0), *pen3-4*, *abcc10-1*, and *abcc10-2* seedlings grown in the presence of the indicated auxins and auxin precursors. Scale bar, 5 mm. **B)** Mean normalized primary root lengths (+SEM; n ≥ 45) of 7-day-old Wt (Col-0), *pen3-4*, *abcc10-1*, and *abcc10-2* seedlings grown in the presence of the indicated auxins and auxin precursors. F-values obtained from two-way ANOVA showed variation, so multiple comparisons using Dunnett’s test was perform with adjusted p-values indicated (∗p ≤ 0.05). **C)** Photograph of representative 7-day-old Wt (Col-0), *pen3-4*, PS200 (*pen3-4 abcc10-*1), *abcc10-*1, *abcc10-2 pen3-4,* and *abcc10-2* seedlings grown in the presence of 1 μM 2,4-DB. Scale bar, 5 mm. **D)** Mean normalized primary root lengths (+SEM; n ≥ 15) of 7-day-old Wt (Col-0), *pen3-4*, PS200 (*pen3-4 abcc10-*1), *abcc10-*1, *abcc10-2 pen3-4,* and *abcc10-2* seedlings grown in the presence of 1 μM 2,4-DB.

To confirm that the defect in *ABCC10* causes *pen3-4* suppression in isolate PS200, we crossed the *ABCC10* insertional allele *abcc10-2* to the *pen3-4* allele of *ABCG36* to generate the *abcc10-2 pen3-4* double mutant. The *abcc10-2 pen3-4* double mutant phenocopied the partial suppression of *pen3-4* hypersensitivity to 2,4-DB seen in isolate PS200 (Fig. 3C and 3D), providing support for a model in which the *ABCC10* defect in PS200 acts as the causative mutation in suppressing *pen3-4* hypersensitivity to the auxin precursor IBA.

### ABCC10 is localized to the tonoplast and expressed in distinct tissues

IBA-derived auxin has been shown to have strong roles in root and shoot development, including regulation of root apical meristem size and cotyledon expansion (reviewed in Frick and Strader, 2018). Additionally, known IBA transporters display specific expression patterns, such as ABCG36/37 localization to the outer lateral domain of root epidermal cells (Strader and Bartel, 2009; Růžička et al., 2010) and TOB1 localization to lateral root initial cells (Michniewicz et al., 2019). Based on these characteristics of known IBA transporters, we examined ABCC10 expression patterns and developmental roles.

ABCC10 was previously identified on the vacuolar membrane (Jaquinod et al., 2007). To confirm ABCC10 localization, we transformed wild type and vac-ck reporter (Nelson et al., 2007) seedlings with a construct driving an N-terminally YFP-tagged genomic copy of *ABCC10* from the strong *35S* promoter from cauliflower mosaic virus. We found that YFP-ABCC10 co-localized with the tonoplast marker (Fig. 4A), consistent with the previous report identifying ABCC10 in tonoplast cell fractions isolated from Arabidopsis cell cultures (Jaquinod et al., 2007; reviewed in Wanke and Üner Kolukisaoglu, 2010). Although we did not detect YFP-ABCC10 signal elsewhere in the cell, we cannot exclude the possibility that a small amount of ABCC10 may localize to other membranes.

**Figure 4.**
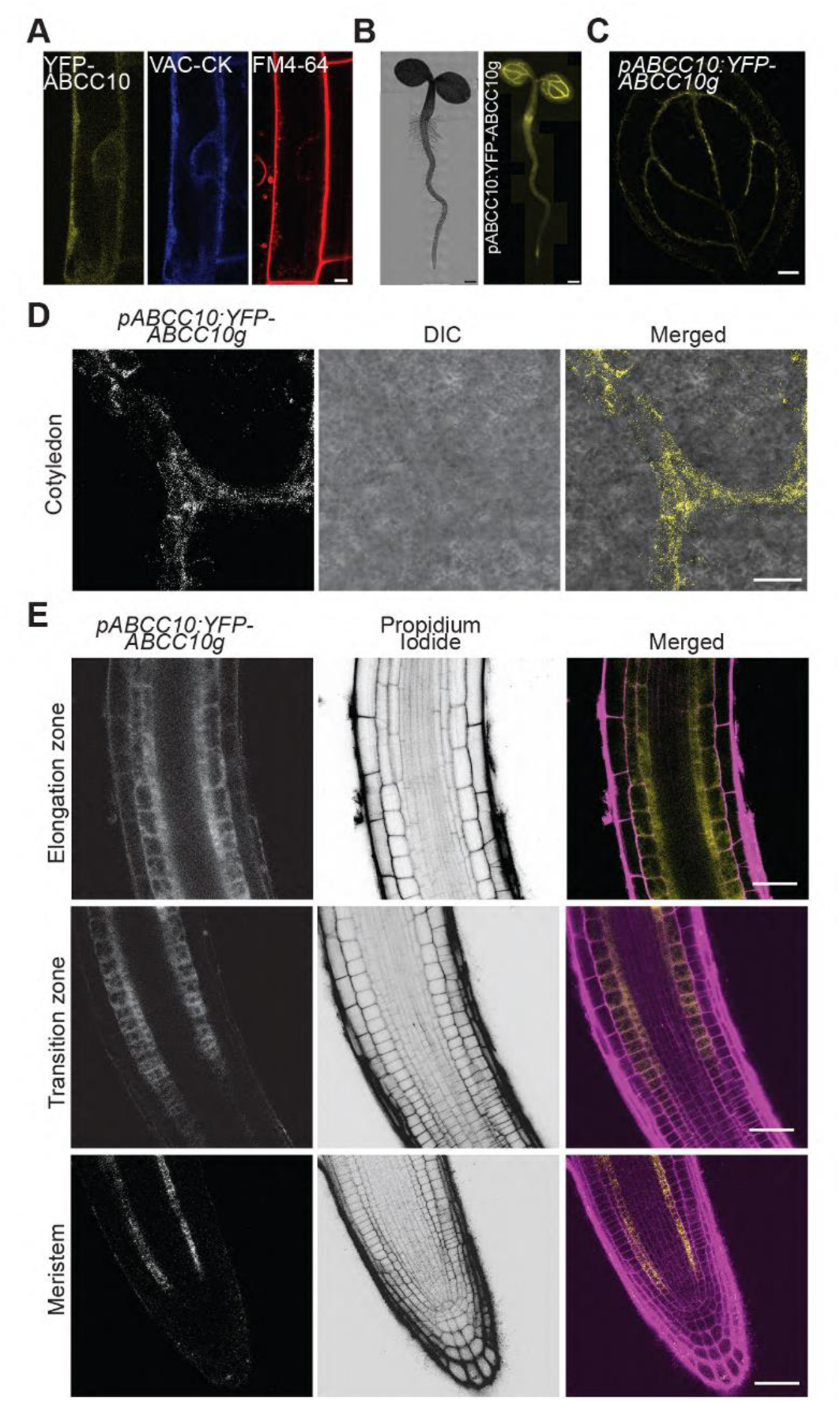
ABCC10 is localized to the vacuolar membrane of the meristematic root cortex, root-shoot junction and the leaf vasculature. **A)** YFP-ABCC10 localizes to the tonoplast in Arabidopsis. Fluorescence microscopy and DIC images from representative 4-day-old Wt (Col-0) co-expressing *p*EG104-ABCC10g and the vac-ck reporter, which decorates the vacuole/tonoplast. Seedlings were stained in 2 μM FM4-64 on ice to label the plasma membrane. Scale bar, 5 μm. **B)** Whole seedling fluorescence microscopy and DIC images of representative 4-day-old Wt (Col-0) and *pABCC10:YFP-ABCC10g* in *abcc10-1* seedlings. Scale bar, 250 μm. **C)** YFP-ABCC10 localizes to the leaf vasculature. Fluorescence microscopy from a representative 4-day-old *pABCC10:YFP-ABCC10g* in *abcc10-1* seedling. Scale bar, 100 μm. **D)** YFP-ABCC10 localizes to the leaf vasculature. Confocal microscopy image of a 4-day-old cotyledon showing expression of *pABCC10:YFP-ABCC10g* in *abcc10-1* seedling in the veins. Scale bar, 100 μm. **E)** YFP-ABCC10 localizes to cortex and endodermis of root. Confocal microscopy of a 4-day-old seedling showing expression of *pABCC10:YFP-ABCC10g* in *abcc10-1* seedling in the cortex and endodermis initials in the root tip (meristematic zone) and in the differentiated cortex and endodermis cells of the elongation zone. Scale bar, 100 μm.

Consistent with published expression databases (Winter et al., 2007; Brady et al., 2007), we found YFP-ABCC10 signal in Arabidopsis seedlings expressing *pABCC10:YFP-ABCC10g* in the leaf vasculature (Fig. 4B-4D), vacuolar membrane of the cortex and endodermis in the root meristem, root elongation zone (Fig. 4E and Supplemental Fig. 1B), and at the root-shoot junction (Supplemental Fig. 1C).We used the ABCC10 expression patterns to guide our examination of *abcc10* mutants for developmental defects.

### *abcc10* mutants display developmental defects in the root apical meristem

The root apical meristem is a small region that undergoes indeterminate growth, generating new root tissue through balanced cell division and differentiation, while maintaining a small group of cells that undergo occasional cell division, called the quiescent center. Maintenance of auxin levels and the establishment of an auxin gradient in this region is essential to proper root patterning and meristem maintenance (reviewed in Iyer-Pascuzzi and Benfey, 2009). IBA-derived auxin regulates root meristem size (Strader et al., 2011) and ABCC10 localizes to the root meristematic zone (Fig. 4B). We therefore examined whether ABCC10 defects affected root meristem development or size. We found no developmental abnormalities in the root meristem of *abcc10* (Fig. 5A); however, the root apical meristem length, measured as the distance between the quiescent center and the first elongating cell of the root, was reduced in *abcc10* compared with wild type (Fig. 5A and 5B), suggesting a role for ABCC10 in regulating root meristem size.

**Figure 5.**
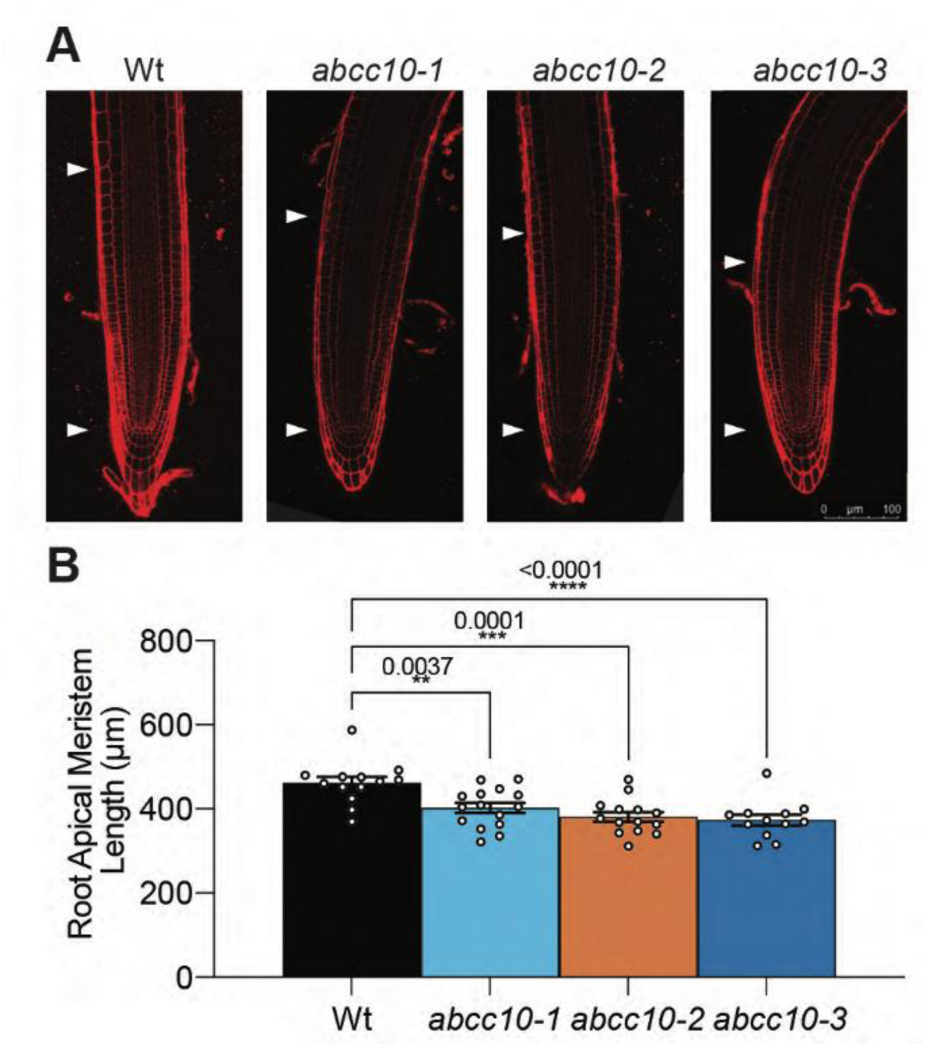
ABCC10 plays a role in regulation of root meristem size. **A)** Fluorescence microscopy of propidium iodide-stained root tips from 4-day-old Wt (Col-0), *abcc10-1, abcc10-2, and abcc10-3*. Arrowheads delineate the top and bottom of the root apical meristem. Scale bar, 100 μm. **B)** Mean root apical meristem lengths (+SEM; n ≥ 15) of 4-day-old Wt (Col-0), *abcc10-1, abcc10-2, and abcc10-3* seedlings. F-values obtained from one-way ANOVA showed variation, so multiple comparisons using Dunnett’s test was perform with adjusted p-values indicated (∗p ≤ 0.05).

### *abcc10* mutants display increased leaf size

Mutants defective in the IBA efflux carrier ABCG36 display hypersensitivity to the inhibitory effects of IBA on cotyledon expansion (Strader and Bartel, 2009). Based on the expression of *ABCC10* in the leaf vasculature of Arabidopsis cotyledons and the weak IBA resistance in roots, we sought to determine whether *abcc10* mutants could suppress the reduced cotyledon size of *pen3-4* in response to IBA or 2,4-DB treatment. Unlike the partial suppression of *pen3-4* hypersensitivity to 2,4-DB in the PS200 isolate observed in root elongation assays (Fig. 1B and 1C), the PS200 isolate (*pen3-4 abcc10-1*) displayed full suppression of the *pen3-4* hypersensitivity to both IBA and 2,4-DB in cotyledon size assays (Fig. 6A and 6B). Furthermore, *abcc10* mutants displayed resistance to the inhibitory effects of IBA and 2,4-DB on cotyledon expansion (Fig. 6A and 6B).

**Figure 6.**
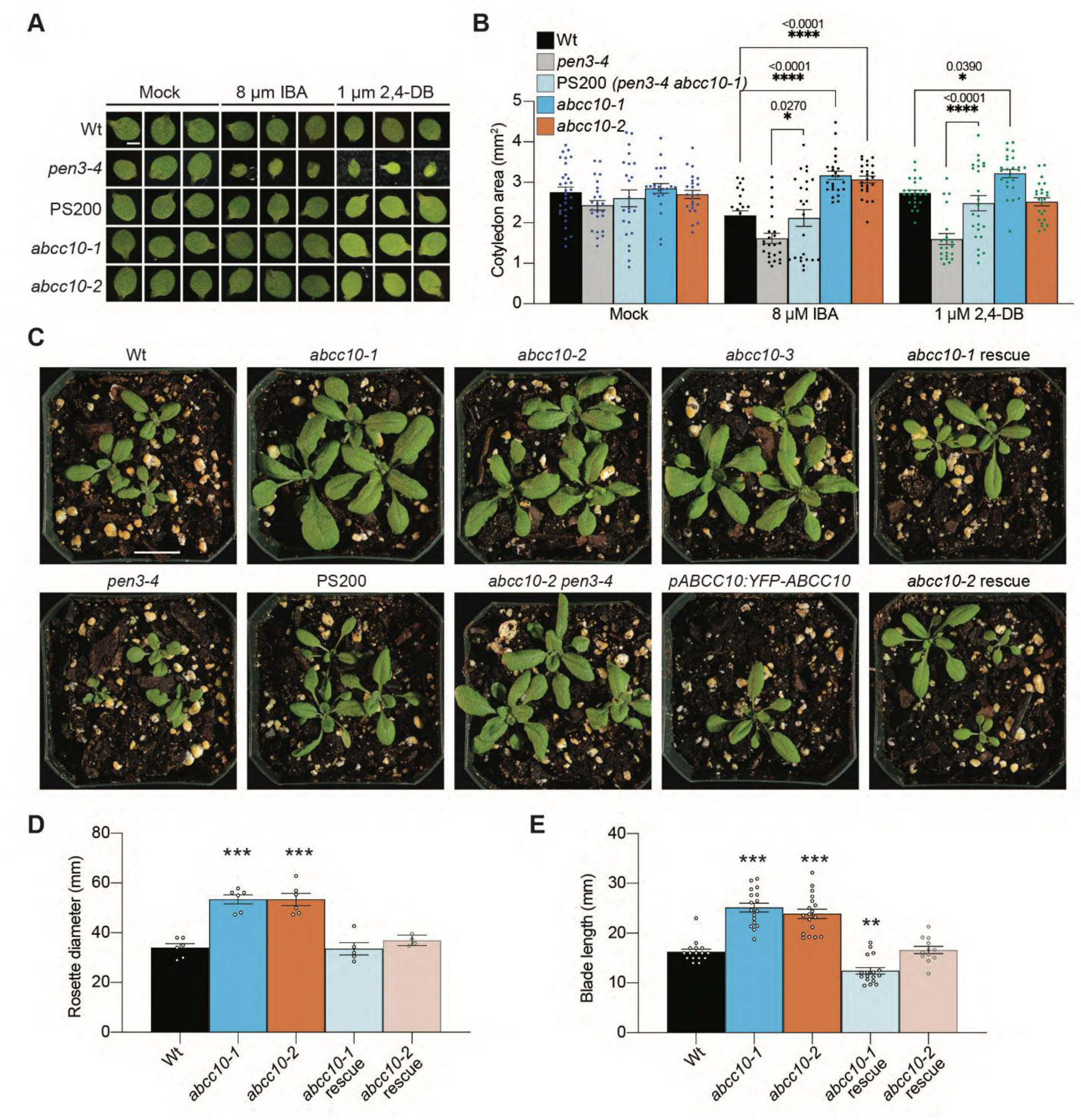
ABCC10 is required for IBA response in cotyledons and regulates leaf development. **A)** Photograph of representative 7-day-old Wt (Col-0), *pen3-4*, PS200, *abcc10-1*, and *abcc10-2* cotyledons from seedlings grown in the presence of the indicated auxins and auxin precursors. Scale bar, 1 mm. **B)** Mean cotyledon area (+SEM; n ≥ 20) of 7-day-old Wt (Col-0), *pen3-4*, PS200, *abcc10-1*, and *abcc10-2* seedlings grown in the presence of the indicated auxins and auxin precursors. F-values obtained from two-way ANOVA showed variation, so multiple comparisons using Dunnett’s test was perform with adjusted p-values indicated (∗p ≤ 0.05). **C)** Soil-grown *abcc10* plants are larger than Wt (Col-0) at 19 d, a phenotype which can be rescued by *pABCC10:YFP-ABCC10g* expression. Scale bar, 20mm. **D)** Mean rosette diameter (+SEM; n ≥ 4) of 19-day-old Wt (Col-0), *abcc10-1*, *abcc10-2*, *abcc10-1* carrying *pABCC10:YFP-ABCC10* (*abcc10-1* rescue), and *abcc10-2* carrying *pABCC10:YFP-ABCC10* (*abcc10-2* rescue) plants. F-values obtained from two-way ANOVA showed variation, so multiple comparisons using Dunnett’s test was perform with adjusted p-values indicated (∗p ≤ 0.05). **E)** Mean leaf blade lengths (+SEM; n ≥ 12) of 19-day-old Wt (Col-0), *abcc10-1*, *abcc10-2*, *abcc10-1* carrying *pABCC10:YFP-ABCC10* (*abcc10-1* rescue), and *abcc10-2* carrying *pABCC10:YFP-ABCC10* (*abcc10-2* rescue) plants. F-values obtained from two-way ANOVA showed variation, so multiple comparisons using Dunnett’s test was perform with adjusted p-values indicated (∗p ≤ 0.05).

IBA-derived auxin has striking effects on cotyledon expansion, suggesting that increased IBA accumulation in cotyledons can increase auxin levels (Strader and Bartel, 2009, 2011; Růžička et al., 2010; Zhang et al., 2016). Although IBA contributions to auxin homeostasis often affects cotyledon expansion in the absence of exogenous auxin, we did not detect any differences in cotyledon size of *abcc10* mutants when compared to wild type (Fig. 6A and 6B).

Although *abcc10* cotyledons (seedling leaves) were of similar size to wild type (Fig. 6A and 6B), adult leaves were markedly larger than those of wild type (Fig. 6C, 6D, and 6E). *abcc10* rosette leaves are larger than those of wild type (Fig. 6C and 6E), consistent with a high-auxin phenotype in this tissue. The larger leaves of *abcc10* can be rescued by expressing *pABCC10:YFP-ABCC10g* in the *abcc10* background (Fig. 6C, 6D, and 6E), confirming a role for ABCC10 in leaf growth.

### ABCC10 transports IBA

The altered IBA and 2,4-DB responsiveness of *abcc10* in roots (Fig. 3A and 3B) and in cotyledons (Fig. 6A and 6B), the identity of *ABCC10* as a member of the MRP subfamily of ABC transporters (Fig. 2D), and *abcc10* suppression of *abcg36* hyperaccumulation of [^3^H]-IBA (Fig. 1E) suggests ABCC10 functions in IBA transport. To test ABCC10 ability to transport IBA, we examined [^3^H]-IBA accumulation in *abcc10* mutants using a modified whole seedling auxin transport assay (Ito and Gray, 2006; Strader et al., 2008; Strader and Bartel, 2009). In this assay, seedlings are incubated for one hour in buffer containing [^3^H]-IBA, followed by scintillation counting. This assay measures the balance of uptake and efflux of [^3^H]-IBA and [^3^H]-IBA-derived molecules (Růžička et al., 2010; Strader et al., 2010) within the assayed tissues. Compared to wild type, *abcc10* seedlings accumulated reduced levels of [^3^H]-IBA (Fig. 7A), suggesting a role for ABCC10 in transport of IBA or IBA-derived molecules in seedlings. In particular, the reduced accumulation of [^3^H]-IBA in *abcc10* (Fig. 7A), combined with its localization to the vacuolar membrane (Jaquinod et al., 2007; reviewed in Wanke and Üner Kolukisaoglu, 2010; Fig. 4A) suggests a role for ABCC10 in moving IBA across the vacuolar membrane.

**Figure 7.**
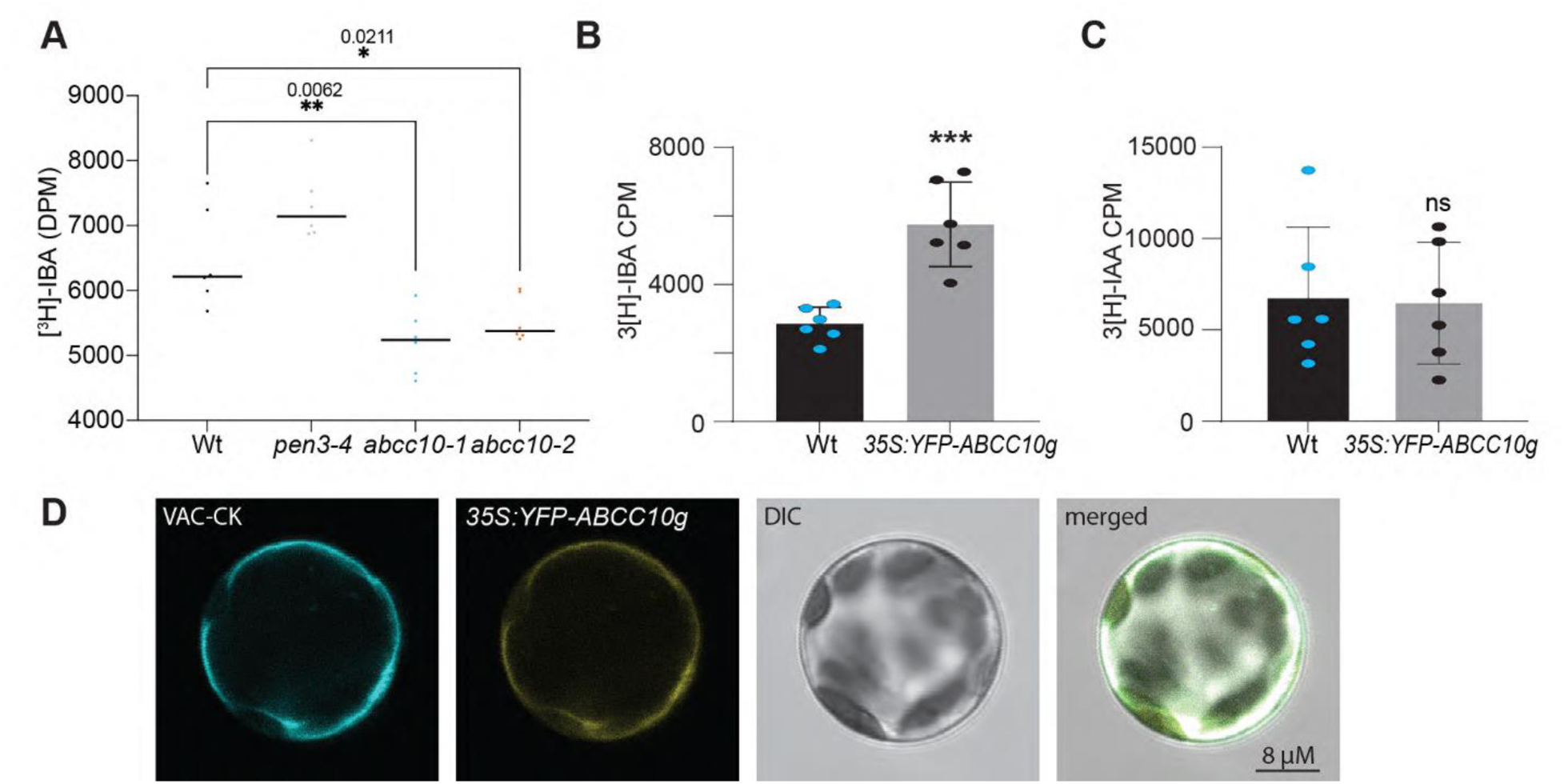
ABCC10 transports IBA. **A)** Whole Wt (Col-0), *pen3-4*, *abcc10-1,* and *abcc10-2* seedlings were incubated for 1 h in uptake buffer containing 25 nM [^3^H]-IBA, rinsed three times with uptake buffer, and then removed and analyzed by scintillation counting. Mean DPM with six replicates of five seedlings of each genotype, are shown. F-values obtained from one-way ANOVA showed variation, so multiple comparisons using Dunnett’s test was perform with adjusted p-values indicated (∗ p ≤ 0.05). **B)** Protoplasts from Wt (Col-0) and Col-0 expressing p*EarleyGate104-ABCC10g* were incubated with 10 nM [^3^H]-IBA. Protoplasts were incubated for 30 min in ice with [^3^H]-IBA and then washed three times before measuring radioactivity. The assay was performed with six biological replicates containing 100,000 protoplasts per replicate and data represents average of six replicates. The F-values obtained from one-way ANOVA showed variation, so multiple comparisons using Dunnett’s test was performed with adjusted p-values indicated (*** p ≤ 0.001). **C)** Protoplasts from Wt (Col-0) and Col-0 expressing p*EarleyGate104-ABCC10g* were incubated with 10 nM [^3^H]-IAA. Protoplasts were incubated for 30 min in ice with [^3^H]-IAA and then washed three times before measuring radioactivity. The assay was performed with six biological replicates containing 100,000 protoplasts per replicate and data represents average of six replicates. The F-values obtained from one-way ANOVA showed variation, so multiple comparisons using Dunnett’s test was performed; no statistical difference was identified (cutoff p ≤ 0.05). **D)** Confocal images of Arabidopsis protoplasts from plants expressing p*EarleyGate104-ABCC10g* and the vac-ck marker showing colocalization of ABCC10 in protoplast vacuoles.

To test whether ABCC10 transports IBA in a simplified system, we examined [^3^H]-IBA or [^3^H]-IAA accumulation in protoplasts generated from wild type (Col-0) or from wild type expressing *35S:YFP-ABCC10* (Fig. 7B and 7C). We found that *YFP-ABCC10*-expressing protoplasts accumulated more [^3^H]-IBA than wild type protoplasts (Fig. 7B). Furthermore, *YFP-ABCC10*-expressing protoplasts accumulated similar levels of [^3^H]-IAA as wild type protoplasts (Fig. 7C). YFP-ABCC10 localizes to the vacuolar membrane in protoplasts (Fig. 7D). This simplified system allows for exclusion of accumulation differences caused by cell-to-cell transport defects; our protoplast data is consistent with a model in which ABCC10 acts to move IBA from the cytoplasm into the vacuole.

## DISCUSSION

IBA-derived IAA controls many aspects of plant growth and development, including root apical meristem size and cotyledon expansion, yet the mechanisms regulating IBA contributions to the active auxin pool remain elusive. Modulation of IBA levels can be achieved on a cellular level to fine-tune IBA contributions to the active auxin pool, as was demonstrated in the work describing the role of TOB1 in vacuolar sequestration of IBA in lateral root initial cells to modulate lateral root formation (Michniewicz et al., 2019). Our identification of ABCC10 as a tonoplast-localized (Fig. 4 and 7) transporter of IBA (Fig. 7) with a role in root apical meristem (Fig. 5) and leaf (Fig. 6) growth provides further insight into the mechanisms by which plants modulate auxin levels to control various aspects of development.

ABCC10 is a member of the MRP/ABCC subfamily of ABC transporters (Fig. 2), which are part of a larger group of ABC transporters in plants (reviewed in Wanke and Üner Kolukisaoglu, 2010). Plant ABC transporters constitute a large and diverse family of primary transporters with a multitude of substrates including hormones, pigments, and secondary metabolites (reviewed in Ha Thi Do et al., 2021). ABC transporters share a common basic structure consisting of a nucleotide binding domain, which contains conserved sequences involved in ATP binding and hydrolysis, and a hydrophobic trans-membrane domain (TMD), involved in translocating the substrate. For many plant ABC proteins, the NBD and TMD domains are fused to generate proteins composed of a single NBD and TMD (“half-size”) or two NBDs and TMDs (“full-size”) (reviewed in Wanke and Üner Kolukisaoglu, 2010). The full-sized ABC proteins, which alone form functional transporters, can be subdivided into three families that were originally named from observations that members of these families confer resistance to various drugs: multidrug resistance (MDR, [TMD–NBD]_2_, ABCB), multidrug resistance-associated protein (MRP, TMD-[TMD–NBD]_2_, ABCC), and pleiotropic drug resistance (PDR, [NBD–TMD]_2_, ABCG) (Crouzet et al., 2006). Further studies into the transport capacities of these proteins have revealed these proteins are capable of transporting a multitude of substrates and have roles other than simply detoxifying cells (reviewed in Ha Thi Do et al., 2021), suggesting these definitions are too restrictive. For instance, many members of the plant MDR/ABCB subfamily have well-characterized roles in IAA transport, whereas some members of the plant PDR/ABCG subfamily have been characterized for their role in IBA transport. Here, we found that MRP/ABCC subfamily member ABCC10 is capable of transporting IBA (Fig. 7).

ABCCs are classically described as glutathione-S conjugate (GSH-X) pumps, due to their ability to transport GSH-conjugates of organic anions (Ishikawa, 1992). The discovery of GSH-X vacuolar import activity was the primary evidence for the existence of ABCC transporters in plants (Martinoia et al., 1993), which were shown to be critical for sequestration and detoxification of compounds to the vacuole (Li et al., 1997). Interestingly, much of the transported IBA is in the form of ester-linked conjugates (Liu et al., 2012) and glutathione deficiency specifically alters lateral and root hair responses to IBA, but not IAA (Trujillo-Hernandez et al., 2020). Despite the apparent role of glutathione in transport and regulation of IBA, it is not clear whether IBA is conjugated to glutathione for transport and/or storage.

The response assays to the inhibitory effects of IBA and 2,4-DB on root elongation (Fig. 1C and 3B) and cotyledon size (Fig. 6B), reduced root apical meristem lengths (Fig. 5), and tonoplast localization (Fig. 4A) of ABCC10 are consistent with IBA efflux from the vacuole. However, our root tip (Fig. 1E) and whole seedling (Fig. 7A) auxin transport assays and the larger rosette leaves in *abcc10* (Fig. 6C and 6D) are consistent with a role for ABCC10 in moving IBA into the vacuole. These alternative models of ABCC10 activity in the cell are each differentially supported by our data; no singular model can explain our current data, suggesting the involvement of additional factors and/or that ABCC10 functions differently in distinct cell types. Despite this limitation, these data clearly demonstrate ABCC10 acts to modulate IBA sensitivity in a variety of contexts, and it is possible that ABCC10 leads to altered IBA contributions to auxin in a tissue- and context-dependent manner. The various phenotypes we have uncovered suggest roles for ABCC10 in both restricting IBA contributions to auxin levels (Fig. 1E, 7A, and 6C) and promoting IBA contributions to auxin levels (Fig. 1C, 3B, 6B, 7B). Future studies will be necessary to resolve the molecular mechanism by which ABCC10 alters IBA contributions to auxin in a cell- and tissue-specific manner.

Our studies have uncovered a previously uncharacterized role for ABCC10 in the regulation of root apical meristem and leaf development through controlling IBA inputs into the auxin pool. Future studies investigating the effect of higher order *abcc* mutants on IBA transport will be instrumental in deciphering the role of this subfamily in auxin homeostasis. The apparent role of glutathione in transport and regulation of IBA, coupled with the preference of ABCCs for GSH-X substrates and the fact that much of long-distance transported IBA is in the form of ester-linked conjugates (Liu et al., 2012), suggest a potential link between ABCC10 and GSH-dependent control of IBA.

Our study further highlights the importance of IBA-derived auxin in various aspects of plant development. Although these results elucidate additional roles for IBA-derived auxin, they demonstrate there remains much left to be uncovered about the molecular mechanism by which IBA-derived auxin contributes to plant growth and development in a cell- and tissue-specific manner.

## MATERIALS AND METHODS

### Plant Growth Conditions

All *Arabidopsis thaliana* lines were in the Columbia (Col-0) background, which was used as the wild type (Wt). Seeds were surface-sterilized (Last and Fink, 1988) and stratified suspended in 0.1% agar for 3d at 4°C to promote consistent germination. Stratified seeds were plated on plant nutrient (PN) media (Haughn and Somerville, 1986) supplemented with 0.5% sucrose (w/v) (PNS) and solidified with 0.6% agar and grown under continuous illumination at 22°C.

### Phenotypic Analyses

To examine auxin-responsive root elongation and cotyledon size, surface-sterilized seeds were stratified for 3d prior to plating on PNS supplemented with hormones or the equivalent amount of ethanol (mock). Treated seedlings were grown horizontally for 7dy under continuous light filtered through a yellow-2208 long-pass plexiglass filter to slow indolic compound breakdown (Stasinopoulos and Hangarter, 1990).

For dark-grown hypocotyl elongation assays, surface-sterilized seeds were stratified for 3d prior to plating on PNS supplemented with hormones or the equivalent amount of ethanol (mock). Treated seedlings were grown horizontally for 1dy under continuous light filtered through a yellow-2208 long-pass plexiglass filter before being wrapped in three layers of aluminum foil to ensure complete blocking of light. The seedlings were then allowed to grow for an additional 5dy before hypocotyls were measured. To examine sucrose dependence, seeds were plated on either unsupplemented PN or PNS and grown horizontally for 1d under continuous white light, followed by growth in the dark for an additional 5dy.

To quantify root apical meristem size, surface-sterilized seeds were stratified for 3d prior to plating on PNS. Seedlings were grown horizontally for 4dy under continuous white light before staining in aqueous propidium iodide (10 μg/mL; Molecular Probes) to label cell walls. Propidium iodide–stained seedlings were imaged using HC PL APO CS2 40x/1.10 WATER lens on a Leica SP8 confocal laser scanning microscope. Samples were excited with the 552-nm laser, and resultant fluorescence between 557 and 784 nm was collected. The distance between the quiescent center and the first elongating cell was measured using FIJI Image software.

To visualize YFP-ABCC10 and γ-TIP-CFP in seedlings, surface-sterilized seeds were stratified for 3d prior to plating on PNS. Seedlings were grown horizontally for 4dy under continuous white light before counterstaining in amphiphilic styryl dye FM4-64 (2 μM; ThermoFisher) on ice to label the plasma membrane (Rigal et al., 2015). FM4-64-stained seedlings were mounted under a coverslip and YFP, CFP, and FM-464 signal from root tips were imaged through a HC PL APO CS2 40x/1.10 WATER lens on a Leica SP8 confocal laser scanning microscope. To detect CFP, samples were excited with the 448-nm laser and the resultant fluorescence between 453 and 509 nm was collected. To detect YFP, samples were excited with the 488-nm laser and the resultant fluorescence between 519 and 547 nm was collected. To detect FM4-64, samples were excited with the 552-nm laser and resultant fluorescence between 557 and 784 nm was collected. The fluorescence intensity of the ROI spline was measured using Plot Profile analysis in FIJI Image Software (Schindelin et al., 2012). The whole seedling image for the plants expressing *pABCC10:YFP-ABCC10g* were acquired using a Leica DMI8 inverted microscope. For imaging the protoplasts of plants expressing both YFP-ABCC10g and γ-TIP-CFP, the protoplasts were prepared using the ‘Tape Arabidopsis sandwich’ method (Wu et al., 2009) and then protoplasts acquired using a Leica DMI8 inverted microscope.

### Vector Construction and transformation

The 5,689 bp *ABCC10* genomic sequence was amplified from Arabidopsis (Col-0) genomic DNA using Pfx platinum (Life Technologies) polymerase in three successive overlapping fragments using primers ABCC10-caccPmlI (5’-CACCACGTGATGATAGAAAATTATTGGACTTCC -3’) and ABCC10-6 (5’-GCTTTCTCCGCCTCTCCCCACCTTG-3’), ABCC10-7 (5’-TGCGGTTCTGTTCTGGTCATCAC-3’) and ABCC10-10 (5’-GGAACAGATACAAACAAGAC-3’), ABCC10-11 (5’-CACACCTCTTGGACGGATTCTTAGCAG-3’) and ABCC10-16 (5’-ATTACTAGTACTACATTTTCCAC-3’), respectively. The first of the ∼2 kb fragments was cloned into the p*ENTR*/D-TOPO vector (Invitrogen), while the other two were cloned into the p*CR4* vector (Life Technologies). Overlap extension PCR cloning (Bryksin and Matsumura, 2010) was then used to link each fragment to obtain p*ENTR-ABCC10g*. This *ABCC10* gDNA was recombined into the p*EarlyGate104* plant expression vector (Earley et al., 2006) using LR Clonase (Invitrogen) to form p*EarleyGate104-ABCC10g* which expresses N-terminal YFP with ABCC10 driven by the cauliflower mosaic virus 35S promoter. The recombinant plasmid was transformed into *Agrobacterium tumefaciens* strain GV3101 (Koncz and Schell, 1986), which was used to transform Col-0 plants using the floral dip method (Clough and Bent, 1998).

Transformants were selected on PN media supplemented with 10 μg/mL Basta (phosphinothricin). Subsequent generations were tested to identify lines homozygous for a single insertion of the transgene, which were then used for downstream experiments.

### Genetic Analysis

Seeds of the *pen3-4* (Stein et al., 2006) mutant were mutagenized by treatment with 0.24% ethyl-methansulfonate (EMS) for 16h (Normanly et al., 1997). The M1 seed were moved to soil and allowed to self-fertilize. The resultant M_2_ seed were surface-sterilized and plated on media supplemented with 10 μM IBA. Seedlings displaying suppressed *pen3-4* hypersensitivity to IBA (Strader and Bartel, 2009) were selected, given a “PS” designation number, moved to soil, and allowed to self-fertilize. M_3_ progeny were retested for suppression of *pen3-4* IBA and 2,4-DB hypersensitivity. Individuals that passed the retest for suppression of *pen3-4* hypersensitivity to the auxin precursors IBA and 2,4-DB were then tested for sensitivity to the active auxins IAA and dichlorophenoxyacetic acid (2,4-D). Isolates displaying resistance to these active auxins were discarded. Remaining isolates displaying a requirement for a fixed carbon source to fuel growth were eliminated because they were likely defective in generic peroxisomal processes (Zolman et al., 2000), rather than processes specific to the auxin precursor IBA. From the remaining isolates, we focused on those displaying suppression of *pen3-4* hyperaccumulation of the [^3^H]-IBA in root tip auxin accumulation assays (Strader and Bartel, 2009). *pen3-4* suppressor isolates PS1, PS88, PS89, PS142, PS173, PS200, and PS219 suppressed the IBA and 2,4-DB hypersensitivity and suppressed the [^3^H]-IBA hyperaccumulation observed in *pen3-4*.

The causative mutation in PS200 was identified by a bulk segregant whole genome sequencing strategy (Thole and Strader, 2015). PS200 was crossed to *pen3-4* and resultant F_2_ progeny were selected for suppression of IBA hypersensitivity, moved to soil, and allowed to self-fertilize. Seedlings from F_3_ progeny that retested for suppression of IBA hypersensitivity were combined for genomic DNA extraction (Thole et al., 2014). Genomic library preparation, Illumina sequencing, and data analysis were performed as previously described (Thole et al., 2014). PCR-based assays were used to verify genotypes. Amplification of *PEN3* with PEN3-Fmut (5’-TCTCTTCTCTGTATCACCCAACTAAATCCTC -3’) and PEN3-Rmut (5’-GAATACCAATCATACCTAAAGCAGACTC -3’) results in a 660-bp product in wild type and no product in *pen3-4*. PCR amplification with PEN3-Fmut and LB1-SALK (5’-CAAACCAGCGTGGACCGCTTGCTGCAACTC -3’) results in an ∼450-bp product in *pen3-4* and no product in wild type. Amplification of *ABCC10* with ABCC10-1 (5’-TGGAGATGAATTAGCAGTTAGGAC -3’) and ABCC10-EcoRV (5’-CGCTCTTCCTTTCGAAGCTCGGGGATAT -3’) results in a 381-bp product with one *EcoR*V in *abcc10-1* and no *EcoR*V site in wild type. PCR amplification with ABCC10-2 (5’-TAACACCTGAGGCTCCTGAAGTAATAGAAG -3’) and ABCC10-12 (5’-GACGTACCTCCCAAATCTCAGCATC -3’) results in a 980-bp product in wild type and no product in *abcc10-2*. PCR amplification with ABCC10-2 and LB1-SALK results in an ∼250-bp product in *abcc10-2* and no product in wild type. PCR amplification with ABCC10-3 F (5’-GATGAACACCGTTACCGTGAGAC -3’) and ABCC10-3 R (5’-AACAGAATCAAAAGCAGGCAAGAAATCCAC -3’) results in a 323-bp product in wild type and no product in *abcc10-3*. PCR amplification with ABCC10-3 F and LB1-SALK results in an ∼200-bp product in *abcc10-3* and no product in wild type.

### Arabidopsis Transport Assays

Auxin accumulation assays were performed as previously described on 5mm excised root tips from 7dy seedlings (Strader and Bartel, 2009) or 4dy whole seedlings, except that root tips or seedlings were moved to scintillant immediately after being rinsed following incubation in [^3^H]-IBA. For the protoplast transport assay, the protoplasts from wild type and plants expressing p*EarleyGate104-ABCC10g* were prepared following the ‘Tape-Arabidopsis Sandwich’ method (Wu et al., 2009). The assay was performed by incubating six biological replicates containing 100,000 protoplasts and 10nM [^3^H]-IBA or [^3^H]-IAA. The protoplasts were incubated for 30 min in ice and then washed three times and then the samples were moved to scintillant for measuring radioactivity.

### Phylogenetic Analysis

Protein sequences corresponding to the ATP-binding cassette, sub-family C domains of ABCC10 and relatives were aligned with Lasergene MegAlign (DNASTAR) using the ClustalW default settings with the Gonnet series protein weight matrix, then manually adjusted to optimize alignments. An unrooted phylogram was generated using by performing the bootstrap method with 500 replicates with distance as the optimality criterion and all characters weighted equally.

### Quantification and Statistical Analysis

Details of statistical analyses can be found in the figure legends. Statistical analysis was performed using Prism 9.1.2 (GraphPad Software, San Diego, CA, United States). All error bars represent standard error of the mean.

## ACCESSION NUMBERS

*ABCC10*, At3g59140; *ABCG36*, At1g59870.

## ACKNOWLEDGEMENTS AND FUNDING

We would like to thank Joseph Cammarata, Hongwei Jing, and Nicholas Morffy for critical comments on this manuscript. This research was supported by the National Science Foundation (IOS-1453750) and the National Institutes of Health (R35GM136338).

## AUTHOR CONTRIBUTIONS

LCS, MM, and LKG performed mutant screen and initial mutant characterization. ALH, SD, and KS confirmed the causative mutation, and performed phenotypic characterization. ALH and SD examined expression patterns and transport activity. ALH wrote the initial manuscript draft with edits by LCS. All authors made edits and approved the final draft. LCS oversaw the project.

---

**Supplementary Figure 1.**
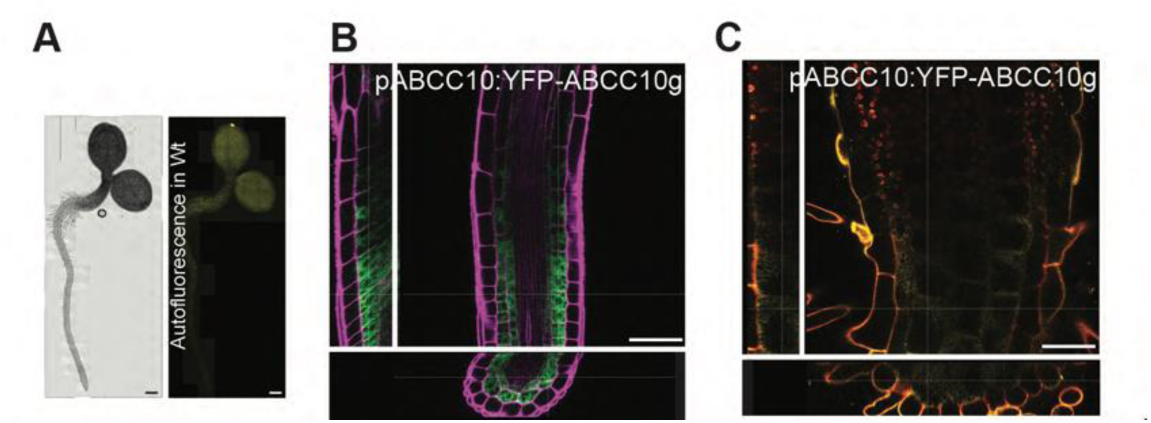
*pABCC10:YFP-ABCC10* is expressed in the root cortex and endodermis **A)** Whole seedling fluorescence microscopy and DIC images of representative 4-day-old Wt (Col-0). Scale bar, 250 μm. **B)** YFP-ABCC10 localizes to the cortex and endodermis of the root meristem. Orthogonal view from different planes (x/y, x/z or y/z) of a fluorescent confocal microscope image from a representative 4-day-old *pABCC10:YFP-ABCC10g* in *abcc10-1* seedling (false-colored green) counterstained with propidium iodide (false-colored magenta). Scale bar, 50 μm. **C)** YFP-ABCC10 localizes to the cortex of the root-shoot junction. Orthogonal view from different planes (x/y, x/z or y/z) of a fluorescent confocal microscope image from a representative 4-day-old *pABCC10:YFP-ABCC10g* in *abcc10-1* seedling (false-colored yellow) counterstained with propidium iodide (false-colored red). Scale bar, 50 μm.

